# Knot or Not? Sequence-Based Identification of Knotted Proteins With Machine Learning

**DOI:** 10.1101/2023.09.06.556468

**Authors:** Denisa Šrámková, Maciej Sikora, Dawid Uchal, Eva Klimentová, Agata P. Perlinska, Mai Lan Nguyen, Marta Korpacz, Roksana Malinowska, Pawel Rubach, Petr Šimeček, Joanna I. Sulkowska

## Abstract

Knotted proteins, although scarce, are crucial structural components of certain protein families, and their roles remain a topic of intense research. Capitalizing on the vast collection of protein structure predictions offered by AlphaFold, this study computationally examines the entire UniProt database to create a robust dataset of knotted and unknotted proteins. Utilizing this dataset, we develop a machine learning model capable of accurately predicting the presence of knots in protein structures solely from their amino acid sequences, with our best-performing model demonstrating a 98.5% overall accuracy. Unveiling the sequence factors that contribute to knot formation, we discover that proteins predicted to be unknotted from known knotted families are typically non-functional fragments missing a significant portion of the knot core. The study further explores the significance of the substrate binding site in knot formation, particularly within the SPOUT protein family. Our findings spotlight the potential of machine learning in enhancing our understanding of protein topology and propose further investigation into the role of knotted structures across other protein families.

**TOC Graphic:** 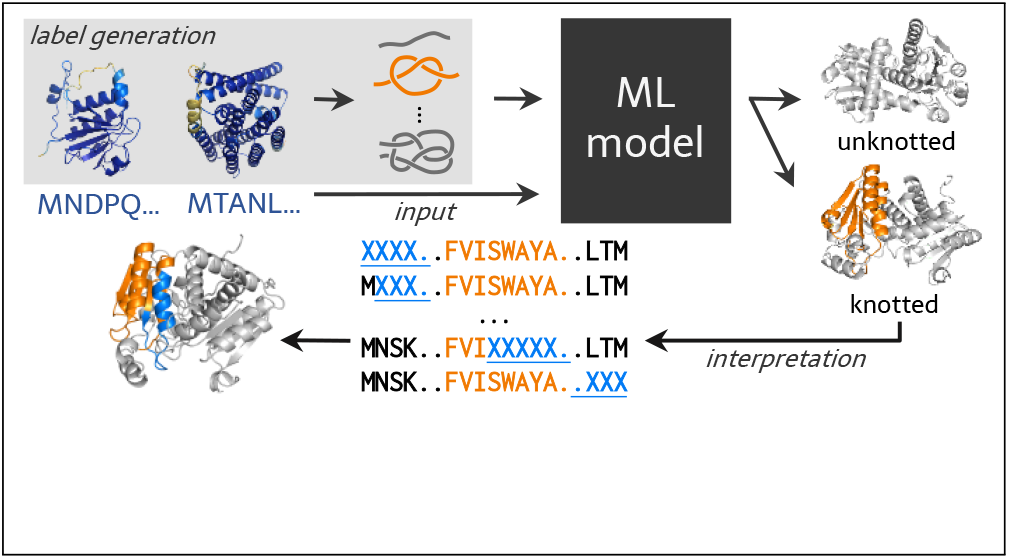

## Introduction

The central paradigm of structural biology is the relationship between protein sequence, the 3D structure it encodes, and biological function. The 3D structure is composed of structural motifs found consistently in unrelated and often sequentially distinct proteins. One type of the motifs whose role is not yet known is knotted motifs, even though they were uncovered three decades ago.^1,2^ The knots are formed by the protein backbone, and the simplest one, with three crossing 3_1_, is shown in Figure 1.

**Figure 1.**
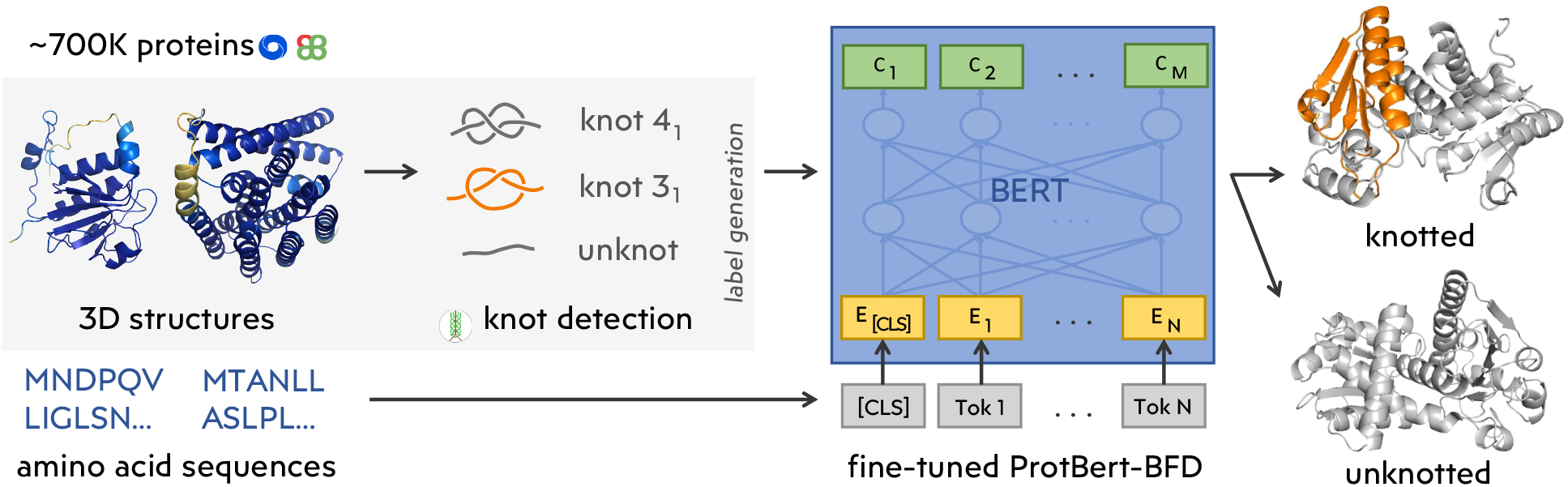
Scheme of the machine learning workflow. The entire approach consists of three main steps: 1. Obtaining data from PDB (with support of Knotprot) and AlphaFold (with support of AlphaKnot) predictions for UniProt; 2. Identifying non-trivial topology in the protein backbone using Topoly; 3. Training a machine learning model to predict non-trivial topology from the protein sequence.

Knots are a common part of our everyday life. In a macromolecular world, they easily form on polymers, however, the analysis of Protein Data Bank shows, that knots are rare in the case of proteins.^3^ Nevertheless, entangled proteins are found in species from all superkingdoms (Bacteria, Eukaryota, and Archaea alike), ^4^ and conduct numerous functions in various parts of the cell.^5,6^ Knots are an integral structural feature of certain protein families^7,8^ where knot forms an active site. Moreover, it is exceptionally conserved, even in families with high sequence diversity. ^5,9,10^ Knotted proteins have already been investigated from different perspectives: folding, ^11–14^ degradation, thermal or mechanical stability^15^ and the potential role of the knot.^8,15^ However, despite broad efforts, the origin, the pattern of amino acids responsible for the entanglement, or the role of the knot is still unknown. ^6^ One of the main reasons for this is the amount of data being too limited for the confident identification of correlations between potential patterns and knotting behavior.

With the update of AlphaFold (AF) database in 2022,^16,17^ the structure predictions for all proteins from the UniProt database (nearly 200 million entries) became available. Therefore, knot detection and identification entered the Big Data age. The predictive quality of AF methods is already widely recognized as many models passed experimental confirmation. ^18,19^ Although knotted proteins account for only 1% of the structures deposited in the PDB, recent studies suggest, that the AlphaFold performs very well here too. AlphaFold has demonstrated accurate prediction of knots in known knotted families including homologs with very low sequence similarity, ^4^ and new type of knots,^20,21^ in particular a composite knot (3_1_#3_1_),^4^ which existence has been experimentally confirmed afterward. ^19^

Herein, we conducted a comprehensive review of all 200 million entries to establish the number of proteins with knotted AlphaFold models. We identified new potentially knotted families and focused especially on those in which different types of topology are predicted. Overall, there are approximately 700,000 knotted proteins (with high-quality prediction – pLDDT value above 70). We classified these proteins into 15 groups (superfamilies), including 10 which members have experimentally determined knotted structures. However, even though it is expected that the knot is strictly conserved in the homologous proteins, we found that each group contains both knotted and unknotted members. Up until now, two types of topology have been observed only in the case of OTC/ATC family. Thus, this data on the one hand seem surprising, on the other allows for the first time to treat knotted and unknotted proteins with Big Data approach and ask some fundamental questions: (i) is it possible to distinguish knotted proteins from unknotted ones based solely on amino acid sequence, (ii) in which families the knot is fully conserved, (iii) does the sequence of a knotted protein contain a pattern responsible for knotting?

Motivated by the significant accomplishments of large protein language models in other fields, herein we utilize our compiled dataset to develop a machine learning model capable of accurately predicting the presence of a knot in a protein from its sequence.

Next, to understand the performance of our machine learning model, we focus on individual families with experimentally verified knotted 3D structures. We ask whether the existence of mixed topology within one family is accurate or if it is a matter of the quality of AF prediction. We investigate four families: SPOUT – the biggest knotted family which possesses 3_1_ and 3_1_#3_1_ knot structures; ^19^ sodium/calcium exchanger integral membrane proteins – a family of transmembrane proteins and deep 3_1_ knot; UCH – big family with 5_2_ knot;^22^ and OTC/ATC family with both knotted (deep 3_1_ knot) and unknotted topologies^23^ confirmed by X-ray structures.

Finally, we apply the machine learning model to seek answers to one fundamental question, undertaken repeatedly in the past, whether there is an amino acid sequence that is responsible for the knotting? The answer may uncover the physical reasons behind the knotting or evolution of the knotted proteins and pave the way to the design of artificial knotted proteins. We specifically focus on the SPOUT superfamily, which holds significant potential for industrial and pharmaceutical applications, such as the design of new antimicrobial drugs.^24^

## Results and discussion

### Dataset creation – identification of all potentially knotted proteins in the Uniprot database based on AlphaFold prediction

In order to investigate what percentage of proteins predicted by AF is potentially knotted, we determined the topology of each structure with a pLDDT greater than 70. The topology was detected using the Topoly package^25^ with the HOMFLY-PT polynomial and with probabilistic closer approach^26,27^ (see Methods section Knot detection in AlphaFold database for details).

We found that knotted proteins occur in less than 0.35% (around 700000) of all sequences deposited in the Uniprot database. Using the CD-HIT clustering method, biological and functional annotation we found that majority of sequences can be classified into 17 superfamilies, shown in the Table 1. Those include families with well known knotted members such as SPOUT, AdoMet synthase, Carbonic anhydrae or UCH families (9 families listed in Table 1);^4,28^ new families with 5_2_ knot OPT oligopeptide transporter (with already confirmed experimentally structures); and families spotted already by others with new type of knots e.g 6_3_ but without experimental verification; ^20,21,29^ and yet new (not spotted before) families such as DUF2254 membrane (IPR018723) or DUF4253 (IPR025349). We also found other knotted families (not listed in Table 1), which include members with more than 3 types of knots in homologous sequences (e.g 0_1_, 3_1_, 5_1_ and 5_2_, they are deposited in the AlphaKnot ^4,21^), very complex knots with 9 crossings (see AlphaKnot) and with less than 700 members. Details about classification into families can be found in the Models and Methods section.

**Table 1:**
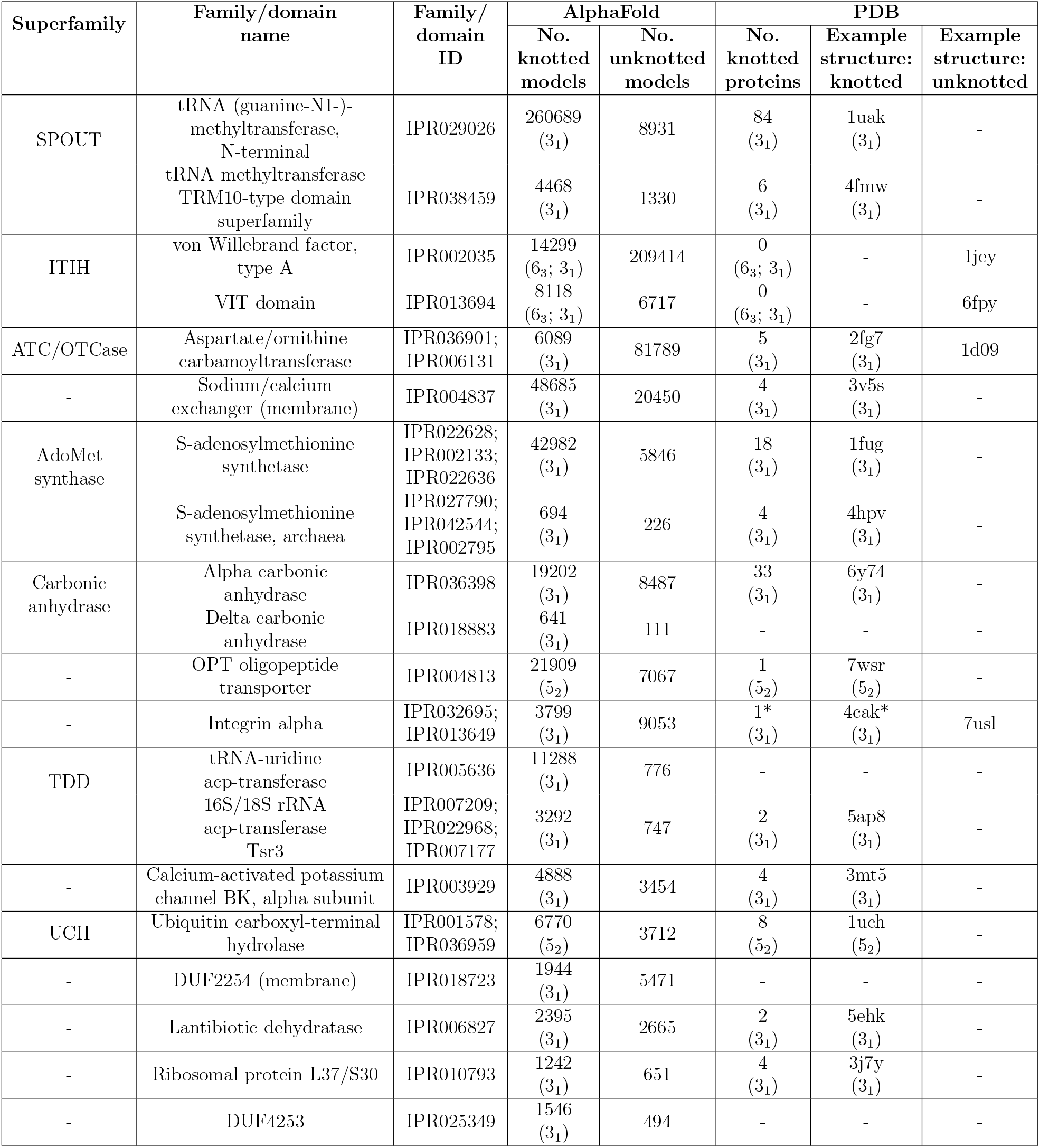
Protein families and domains of the dataset. In parentheses are the types of knots found in PDB structures and AlphaFold models. In the case of models, the dominant knot types are given. In the case of knotted proteins with PDB structures, the number of proteins with topology specified in parentheses is provided. Asterisks indicate that the topology is uncertain due to gaps in key regions of the structures.

| Superfamily | Family/domain name | Family/domain ID | AlphaFold |  | PDB |  |  |
| --- | --- | --- | --- | --- | --- | --- | --- |
|  |  |  | No. knotted models | No. unknotted models | No. knotted proteins | Example structure: knotted | Example structure: unknotted |
| SPOUT | tRNA (guanine-N1-)-methyltransferase, N-terminal | IPR029026 | 260689<br>(3<sub>1</sub>) | 8931 | 84<br>(3<sub>1</sub>) | 1uak<br>(3<sub>1</sub>) | - |
|  | tRNA methyltransferase TRM10-type domain superfamily | IPR038459 | 4468<br>(3<sub>1</sub>) | 1330 | 6<br>(3<sub>1</sub>) | 4fmw<br>(3<sub>1</sub>) | - |
| ITIH | von Willebrand factor, type A | IPR002035 | 14299<br>(6<sub>3</sub>; 3<sub>1</sub>) | 209414 | 0<br>(6<sub>3</sub>; 3<sub>1</sub>) | - | 1jey |
|  | VIT domain | IPR013694 | 8118<br>(6<sub>3</sub>; 3<sub>1</sub>) | 6717 | 0<br>(6<sub>3</sub>; 3<sub>1</sub>) | - | 6fpy |
| ATC/OTCase | Aspartate/ornithine carbamoyltransferase | IPR036901;<br>IPR006131 | 6089<br>(3<sub>1</sub>) | 81789 | 5<br>(3<sub>1</sub>) | 2fg7<br>(3<sub>1</sub>) | 1d09 |
| - | Sodium/calcium exchanger (membrane) | IPR004837 | 48685<br>(3<sub>1</sub>) | 20450 | 4<br>(3<sub>1</sub>) | 3v5s<br>(3<sub>1</sub>) | - |
| AdoMet synthase | S-adenosylmethionine synthetase | IPR022628;<br>IPR002133;<br>IPR022636 | 42982<br>(3<sub>1</sub>) | 5846 | 18<br>(3<sub>1</sub>) | 1fug<br>(3<sub>1</sub>) | - |
|  | S-adenosylmethionine synthetase, archaea | IPR027790;<br>IPR042544;<br>IPR002795 | 694<br>(3<sub>1</sub>) | 226 | 4<br>(3<sub>1</sub>) | 4hvp<br>(3<sub>1</sub>) | - |
| Carbonic anhydrase | Alpha carbonic anhydrase | IPR036398 | 19202<br>(3<sub>1</sub>) | 8487 | 33<br>(3<sub>1</sub>) | 6y74<br>(3<sub>1</sub>) | - |
|  | Delta carbonic anhydrase | IPR018883 | 641<br>(3<sub>1</sub>) | 111 | - | - | - |
| - | OPT oligopeptide transporter | IPR004813 | 21909<br>(5<sub>2</sub>) | 7067 | 1<br>(5<sub>2</sub>) | 7wsr<br>(5<sub>2</sub>) | - |
| - | Integrin alpha | IPR032695;<br>IPR013649 | 3799<br>(3<sub>1</sub>) | 9053 | 1*<br>(3<sub>1</sub>) | 4cak*<br>(3<sub>1</sub>) | 7usl |
| TDD | tRNA-uridine acp-transferase | IPR005636 | 11288<br>(3<sub>1</sub>) | 776 | - | - | - |
|  | 16S/18S rRNA acp-transferase Tsr3 | IPR007209;<br>IPR022968;<br>IPR007177 | 3292<br>(3<sub>1</sub>) | 747 | 2<br>(3<sub>1</sub>) | 5ap8<br>(3<sub>1</sub>) | - |
| - | Calcium-activated potassium channel BK, alpha subunit | IPR003929 | 4888<br>(3<sub>1</sub>) | 3454 | 4<br>(3<sub>1</sub>) | 3mt5<br>(3<sub>1</sub>) | - |
| UCH | Ubiquitin carboxyl-terminal hydrolase | IPR001578;<br>IPR036959 | 6770<br>(5<sub>2</sub>) | 3712 | 8<br>(5<sub>2</sub>) | 1uch<br>(5<sub>2</sub>) | - |
| - | DUF2254 (membrane) | IPR018723 | 1944<br>(3<sub>1</sub>) | 5471 | - | - | - |
| - | Lantibiotic dehydratase | IPR006827 | 2395<br>(3<sub>1</sub>) | 2665 | 2<br>(3<sub>1</sub>) | 5ehk<br>(3<sub>1</sub>) | - |
| - | Ribosomal protein L37/S30 | IPR010793 | 1242<br>(3<sub>1</sub>) | 651 | 4<br>(3<sub>1</sub>) | 3j7y<br>(3<sub>1</sub>) | - |
| - | DUF4253 | IPR025349 | 1546<br>(3<sub>1</sub>) | 494 | - | - | - |

An intriguing observation is that each family contained knotted and unknotted structures, what was only observed in the case of ATCase/OTCase family. In this case, AF predicts only 11% of knotted proteins. The most unexpected are cases of families where the knot was expected to be strictly conserved such as SPOUT (7% unknotted), transmembrane proteins (13% unknotted), or UCH (26% unknotted). Later on, we will show that in some cases we can explain these surprising results.

The data from Table 1 presents a great deal of information that can be analyzed in a variety of ways, however, because it includes families with knotted and unknotted members (with high pLDDT) it presents an ideal dataset for the application of machine learning. Moreover, we will focus on the biggest families, with not more than two types of topology, so families with 4_1_, and 6_1_ knot types have been omitted here. Data selected for ML application are presented in Table 2.

**Table 2:** Evaluation of the finetuned ProtBert-BFD model on the test set. Accuracy, true positive rate (TPR) and true negative rate (TNR) of the model evaluated for the whole test set and per protein family.

| Protein superfamily | Dataset size | Unknotted part size | Accuracy (%) | TPR (%) | TNR (%) |
| --- | --- | --- | --- | --- | --- |
| All | 39412 | 19718 | 98.5 | 98.7 | 98.3 |
| SPOUT | 7371 | 550 | 98.9 | 99.5 | 90.9 |
| ITIH | 14263 | 12555 | 98.7 | 94.2 | 99.3 |
| ATCase/OTCase | 3799 | 3352 | 100.0 | 99.8 | 100.0 |
| Sodium/calcium exchanger | 5256 | 726 | 99.0 | 99.7 | 95.3 |
| AdoMet synthase | 1794 | 240 | 99.0 | 99.3 | 97.1 |
| Carbonic anhydrase | 1531 | 539 | 95.9 | 97.4 | 93.1 |
| OPT Oligopeptide transporter | 2510 | 456 | 98.7 | 99.5 | 94.7 |
| Integrin alpha | 332 | 224 | 83.1 | 66.7 | 91.1 |
| TDD | 612 | 24 | 99.0 | 99.7 | 83.3 |
| Calcium-activated potassium channel BK, alpha subunit | 127 | 87 | 89.0 | 97.5 | 85.1 |
| UCH | 477 | 125 | 90.6 | 96.0 | 75.2 |
| DUF2254 membrane | 593 | 376 | 99.8 | 99.5 | 100.0 |
| Lantibiotic dehydratase | 392 | 286 | 96.4 | 95.3 | 96.9 |
| Ribosomal protein L37/S30 | 147 | 41 | 85.7 | 100.0 | 48.8 |
| DUF4253 | 123 | 53 | 86.2 | 90.0 | 81.1 |

### Machine learning classifies proteins as knotted or unknotted

Next, we ask if there is a machine learning model capable of solving the binary classification of proteins into knotted and unknotted classes based solely on their amino acid sequences. This inquiry was based on the dataset, encompassing 39412 entries, that we constructed by determining the topology of each structure predicted by AlphaFold (further details are outlined in the Models and Methods section). The dataset consists of knotted and unknotted amino acid sequences from 20 different protein families with various knot types (with the prevalence of 3_1_ knot), the details about the number of representants from each family can be found in Table 1. Combining the dataset from individual protein families implies that the dataset is diverse. Sequences within one family are typically similar to each other, but diverse from proteins from other families and each family usually has a different biological function. The sequences also show differences across families in features such as average sequence length or average knot core size. During the evaluation, we thus approach the dataset from two perspectives: the first is a Big Data approach, where we treat the data as a whole and are testing the performance of the model on a big scale, and the second is a more detailed analysis and interpretation of results for individual selected families.

We made the decision to train three distinct models for the task of knot detection, where each model incorporated a progressively less complex architecture and strategy. This approach allowed us to compare the performance of the models and select the most optimal one. Furthermore, it enabled us to investigate whether solving the problem necessitates the use of a large, complex model or if a simpler approach suffices. The first model was a fine-tuned ProtBert-BFD^30^ network, the second was a convolutional neural network using ProtBert-BFD embeddings as input, and the third was a simple convolutional neural network that operated on sequences with one-hot encoded amino acids (further details can be found in the Models and Methods section). Refer to Figure 1 for a summary of the dataset creation process and model training.

After applying machine learning to our dataset, we found that it is possible to distinguish knotted from unknotted proteins based solely on their amino acid sequence. All models showed robust results on the test set (accuracy above 95%), with the first model, the fine-tuned ProtBert-BFD model, demonstrating the best overall accuracy of 98.5%. As our dataset consists of a collection of distinct protein families, where proteins from different families differ in various respects, assessment of models across individual families is necessary. After the evaluation, we chose the first fine-tuned ProtBert-BFD model for subsequent analysis as it displayed strong accuracies across all the different protein families (see Table 2). We attribute the superior performance of the final model to the usage of a pre-trained protein language model. As we can see on its visualized t-SNE embeddings in the Supplementary Figure 1, the model already contained some information about the protein knotting status even before fine-tuning it for the knotting task. In the following sections, we will focus on individual protein families and delve into a qualitative examination of the chosen model.

### Unknotted proteins from knotted families are fragments

Our dataset primarily consists of protein families with experimentally verified knots. However, even within these known to be knotted families, there are cases of unknotted structures. This is quite unexpected since the only one example of a mixed topology within a single family reported until now was the ATC/OTC family^23^ (which is also part of our dataset, see the section below). Therefore, we examined such unknotted cases to determine whether they are truly unknotted or simply a result of database inaccuracies, such as AlphaFold’s incorrect structure prediction or fragmented sequences in UniProt.

We first analyzed the SPOUT superfamily, the largest knotted family in our dataset with known 3D structures but only with a knot. In this case AF predicts 7% of unknotted proteins. We examined the correctness of these structures by analyzing their homologs: each protein was sequentially aligned with its closest knotted homolog to determine whether it contained a region corresponding to the knot core.

The analysis revealed that all unknotted proteins from the SPOUT superfamily have a close homolog (with an average sequence identity of 80%) that is knotted. These proteins are merely fragments of their homologs and lack at least part of the knot core, hence they cannot form knots. Predominantly, they lack the C-terminal part of the knot core, but in some instances, they also miss the N-terminal part or even the entire knotted region (Figure 2). These proteins would not be functional without a fully manifested knot core, as this is an essential component of the SPOUT active site. Given that more than 75% of these structures are annotated as fragments, we conclude that the SPOUT superfamily doesn’t contain any unknotted functional methyltransferases.

**Figure 2.**
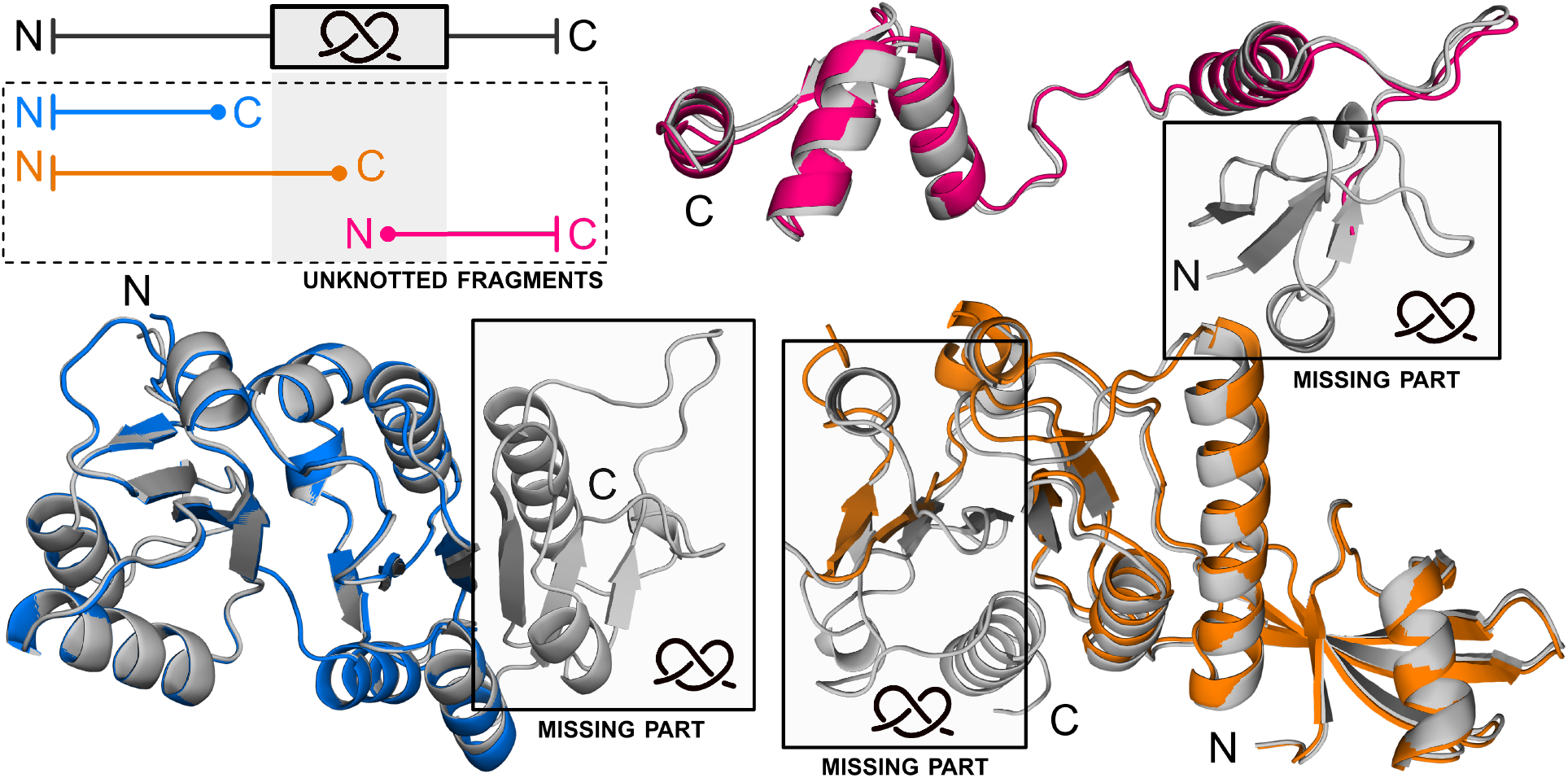
Unknotted SPOUT proteins superposed with their knotted homologs. Upper part: unknotted structure (magenta; UniProtKB ID: A0A418H8R5) lacking the N-terminal part of the knot core. Lower part: unknotted structure (blue; UniProtKB ID: A0A090WL68) without the entire knot core, and unknotted structure (orange; UniProtKB ID: A0A356CS99) without the C-terminal part of the knot core. All knotted homologs are shown with grey color and their knotted regions are marked (UniProtKB IDs: A0A2X1NEC8, A0A090W861, and A0A3D0W667, respectively).

A similar trend was observed in the UCH superfamily. In the case of this superfamily, 17.4% of the proteins in our test set are unknotted even though a strict conservation of knot between several proteins despite their large sequence divergence has been shown. ^5^ Investigation revealed that all unknotted proteins have a close, knotted homolog (with an average sequence identity of 80.5%), and 92% of them are annotated as fragments (Figure 3). Like in the SPOUT members, the unknotted proteins lack at least a portion of the knot core, rendering them unknotted. Interestingly, we noticed that in some rare cases, eukaryotic proteins may exhibit topologically distinct isoforms due to alternative splicing. ^31^

**Figure 3.**
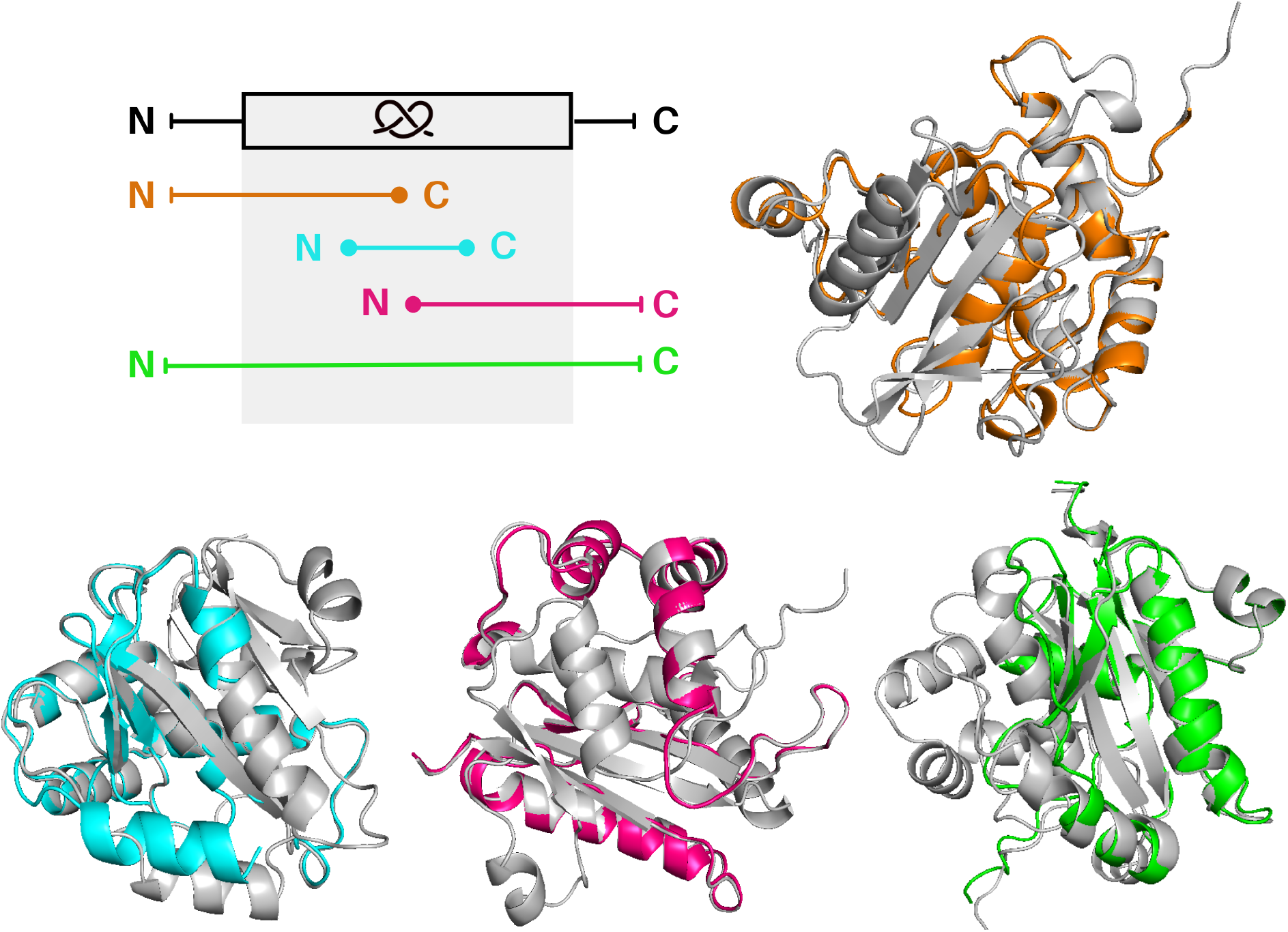
Unknotted UCH proteins superposed with their knotted homologs. Upper part: unknotted structure (orange; UniProtKB ID: A0A7S3I8E7) lacking the C-terminal part of the knot core. Lower part: unknotted structure (cyan; UniProtKB ID: Q06AV7) without the N-terminal and C-terminal part of the knot core, unknotted structure (magenta; UniProtKB ID: A0A1A7YJ99) without the N-terminal part of the knot core, and unknotted structure (green; UniProtKB ID: A0A024WZA) with low quality part of the knot core. All knotted homologs are shown with grey color (UniProtKB IDs: A0A078B145, A0A6P6PV05, A0A1A7X9F0 and W7F0M6, respectively).

The same results were obtained also for the family of transmembrane proteins (see Figure 4). This is the only family consisting of knotted membrane proteins (sodium/calcium exchanger integral membrane proteins),^28^ and here also a stricl conservation of a knot topology was suggested based on the available 3D structures. ^32^ All unknotted proteins have a close homolog that is knotted (with, on average, 79,5% sequence identity). Based on their pairwise alignment, most of the unknotted sequences either align with a fragment starting before the knot core and ending on a residue inside it, or with a fragment starting inside the knot core and ending after it. Thus, due to the lack of a knot core sequence, these proteins cannot form the non-trivial topology. In this case, 56% of the unknotted proteins are annotated as fragments.

**Figure 4.**
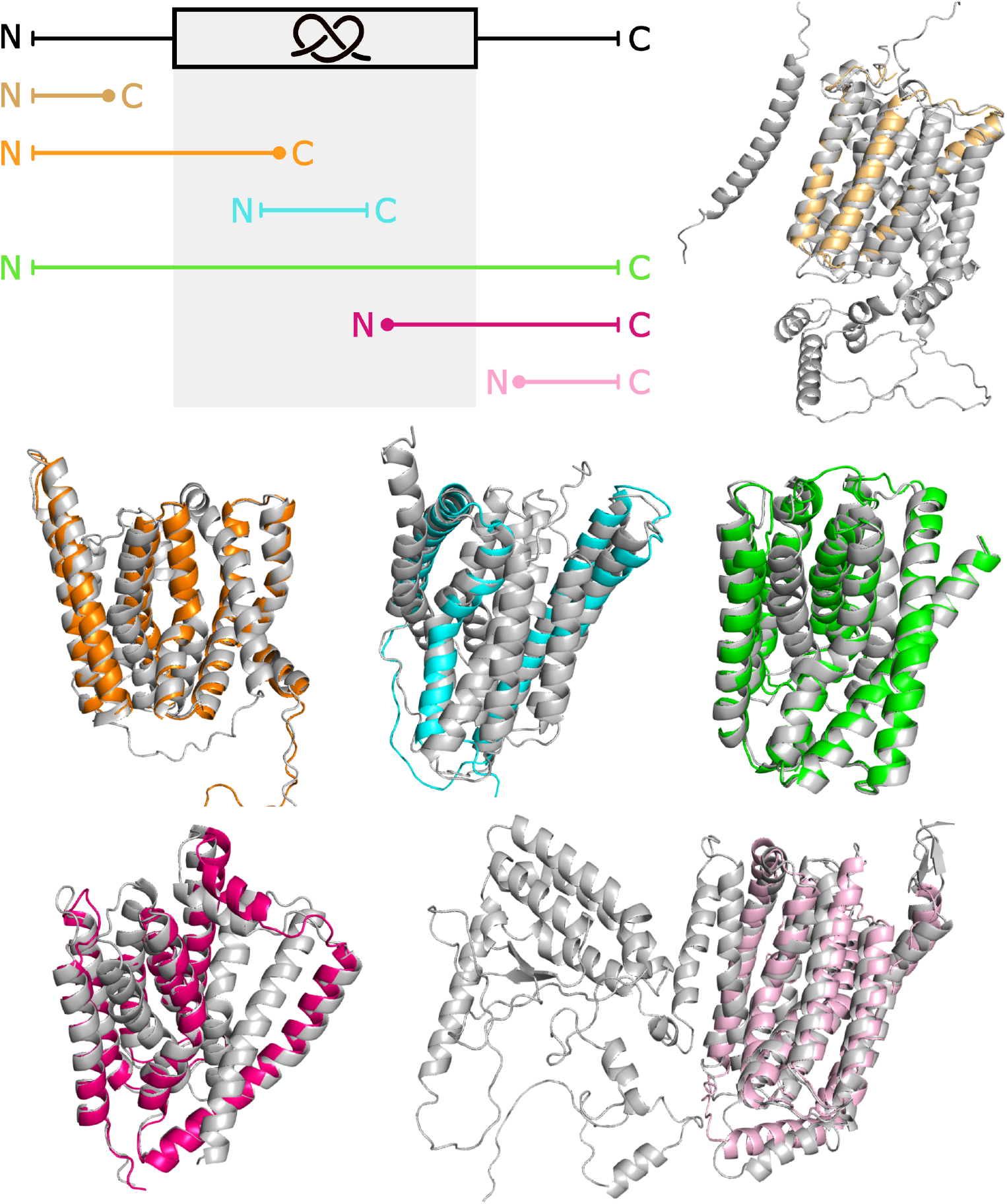
Unknotted membrane proteins from PF01699 family superposed with their knotted homologs. Upper part: unknotted structure (light orange; UniProtKB ID: A0A5K0X7L3) without the entire knot core, unknotted structure (orange; UniProtKB ID: A0A397EWI1) structure lacking the C-terminal part of the knot core, and unknotted structure (green; UniProtKB ID: A0A562DKV6) with low quality part of the knot core. Lower part: unknotted structure (cyan; UniProtKB ID: A0A453N937) without the N-terminal and C-terminal part of the knot core, unknotted structure (magenta; UniProtKB ID: A0A378FTV0) without the N-terminal part of the knot core, and unknotted structure (light pink; UniProtKB ID: K1RUG8) without the entire knot core. All knotted homologs are grey with their knot cores colored in light blue (UniProtKB IDs: V4U0P6, A0A397A8U0, E6WVS2, A0A453N959, A0A0E1C8E8, A0A0L8I7H8, respectively).

Notably, all these families have experimentally verified knotted proteins, while the unknotted ones are based on AlphaFold’s predicted models. By examining these models, we have shown that there are no unknotted proteins within established families of knotted proteins and that their topology is conserved.

### ATC/OTC – a family with both knotted and unknotted proteins

The only family currently known experimentally to possess both knotted and unknotted proteins is ATC/OTC. More specifically, within this family, several subfamilies with a conserved topology are indicated: the unknotted subfamilies include ATC, OTC and PTC, while the knotted subfamilies involve AOTC, SOTC and YTC.^33^ To ascertain whether the proteins with a given topology in our set belong to the appropriate subfamilies (i.e., whether the topology of the subfamilies is conserved), we clustered their sequences and aligned them to assign the closest homologs. This analysis showed that there are no topologically misclassified proteins from this family in our dataset – all unknotted proteins have closest homologs from unknotted subfamilies, and all knotted proteins from knotted subfamilies. Thus, our model can accurately differentiate between topologically different members of the ATC/OTC family.

### Substrate binding site important for knot detection

In order to determine what specific regions of proteins play a role in knot formation, we assessed the significance of continuous parts of the protein sequences (referred to as “patches”) using the model to obtain the information about their importance on the prediction. This process is explained in detail in the Models and Methods section. To ensure the analysis was conducted on high-quality data, we focused on the most significant patches from the largest group of knotted proteins, the SPOUT superfamily. Given that the average knot size in these proteins is about 47 amino acids, we chose patch lengths of 10, 20, and 40 residues. Our results indicate that longer patches, on average, hold more significance (Table 3).

**Table 3:** Percentage of proteins with a patch located in the knot core. Only patches that have at least 50% of their length inside the knot core were calculated.

| Patch size | Significant | Not significant |
| --- | --- | --- |
| 10 | 55% | 24% |
| 20 | 84% | 7% |
| 40 | 95% | 2% |

We started the analysis by localizing the positions of these patches in the protein sequences. This revealed a significant over-representation in a specific region – the knot core. For instance, more than 80% of the patches of length 20 are found within the knotted region. Considering that the majority of these regions contain less than 50 amino acids, and the proteins are on average 250 amino acids long, this accumulation within a specific region suggests that this region is crucial for knot formation, confirming the robustness of our method. Additionally, regardless of their length, all patches point to the same location within the knot – its C-terminal part (Figure 5B). Based on the multiple sequence alignment of the knot cores, we observed that this region is the most conserved part of the knot, containing the glycine motif responsible for substrate binding (Figure 5C). This motif is also present in patches of lower significance, especially those of length 10. However, the conserved glycines are not a unique feature of knotted methyltransferases. They are also found in proteins with other folds, such as Rossmann fold, which are unknotted. ^34^

**Figure 5.**
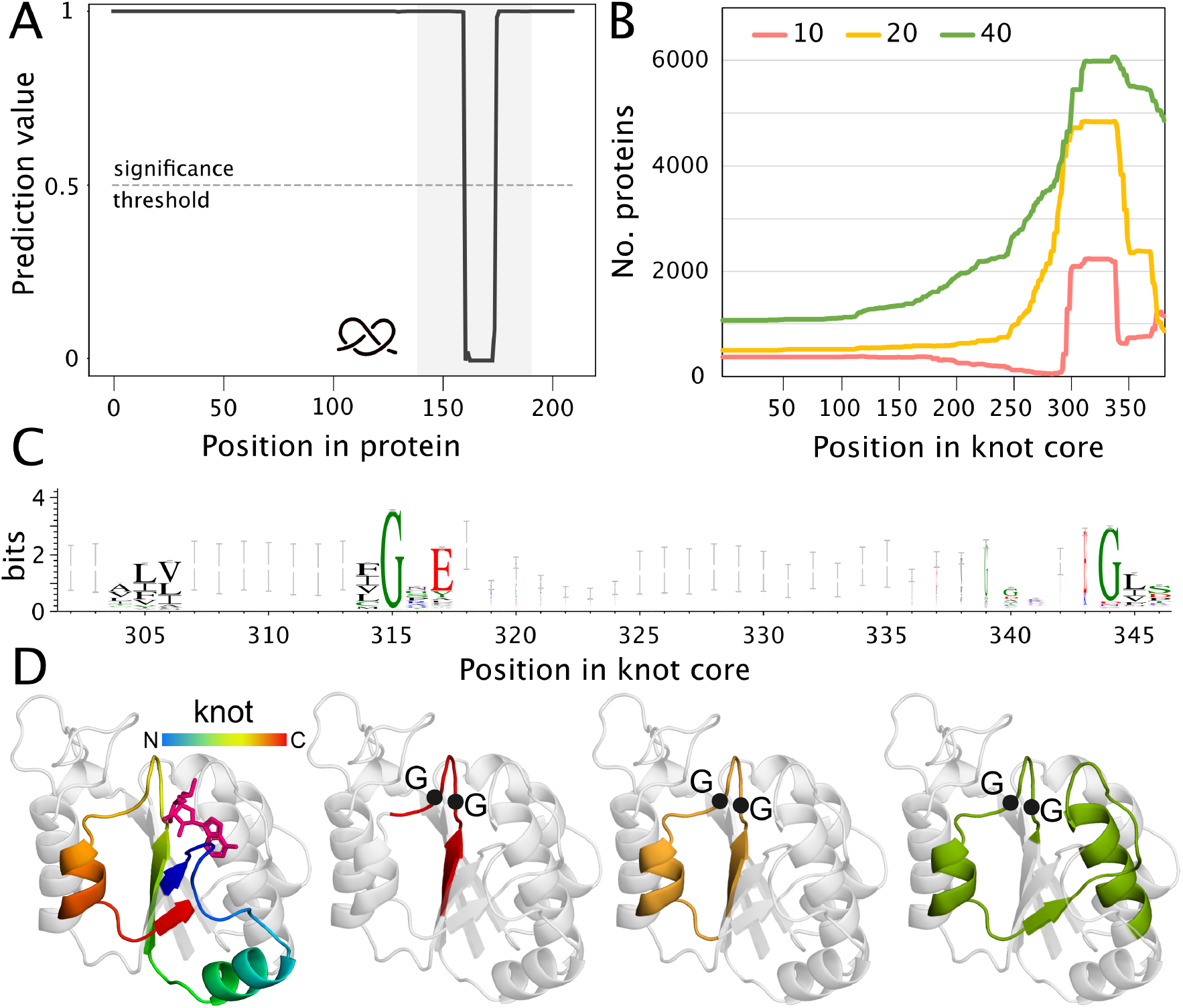
Important regions in SPOUT proteins based on ML model. A. An example of how the model’s prediction changes with respect to the patch location based on Nep1 protein (patches of length 10; UniProtKB ID: Q57977). Note the one significant drop inside the knot core. B. Cumulative positions of the patches (with different lengths: 10, 20, and 40 amino acids) based on the knot core alignment. C. Logo of the C-terminal part of the knot core showing the conserved glycine motif. D. Positions of patches in a TrmD structure (based on N-terminal domain of AlphaFold model of UniProtKB ID: B6YQ31). Colored regions represent from the left: knotted region (rainbow) and estimate location of the ligand (magenta); 10 (red), 20 (yellow) and 40 (green) amino acid-long patches, respectively. Black dots show the position of the conserved glycine residues.

## Discussion

In this study, we have assembled the largest dataset of knotted and unknotted proteins by analyzing AlphaFold’s protein structure predictions using Topoly. Built upon this dataset, we have presented a novel transformer-based machine learning model able to predict the knotting status only from the amino acid protein sequence. The model demonstrated robust performance across different protein families, indicating its effectiveness in predicting the presence of knots on a large scale. However, the training scheme with balanced dataset classes limits the model’s performance on tasks such as scanning the whole UniProt database, where it can behave more like a sensitive sifter for potential knots requiring additional analysis. The performance of the model outside its domain, i.e. on other protein families, is not guaranteed, the results for one such family (ketol-acid reductoisomerases) can be found in Supplementary Table 1.

To facilitate easy access to our model, we provide the link to the Hugging Face hub (https://huggingface.co/roa7n/knots_protbertBFD_alphafold) containing the trained model, along with scripts for creating the dataset, model training and interpretation (https://github.com/ML-Bioinfo-CEITEC/pknots_experiments).

Subsequent utilization of the proposed interpretation patching technique allowed for a deeper analysis of individual proteins and a precise assessment of patterns associated with the presence of knots in those proteins. The patching analysis conducted on the SPOUT protein family marked as important for knotting areas of the co-factor binding site, essential for the family representatives’ enzymatic function. Due to the complexity of the interpretation analysis, our examination was restricted to a specific protein family, leaving room for future investigation of additional sequence patterns in other families, thereby enhancing our understanding of the origin of protein knots. Potential extension to the interpretation would be the usage of different techniques able to interpret longer pieces of the sequence such as modifying the SHAP values.^35^

Another significant finding of this study is the observation that unknotted proteins in our dataset coming from known knotted families are typically non-functional fragments. Even though a significant portion of this data is not annotated as fragments we show in the example of the SPOUT superfamily, UCH superfamily, and transmembrane protein family that unknotted proteins in these families lack at least a portion of the knot core and are thus nonfunctional. Even though the unknotted fragments are valuable and important for the Big Data approach in the machine learning part of this paper, the incompleteness of the crucial domain necessary for the functionality diminishes their biological significance. Additionally, we conducted an analysis of the ATC/OTC family, which is currently the only known family with mixed knotted and unknotted topology. Our investigation showed that a given topology is conserved within individual subfamilies. However, we managed to examine only a small part of the families containing proteins with knot, leaving thus space for future investigation of other knotted protein families. Those unexplored families may either follow the scheme of SPOUT, UCH and transmembrane proteins and be fully knotted with unknotted proteins being only non-functional fragments, or there might exist another family similar to ATC/OTC with both knotted and unknotted fully functional proteins.

## Conclusions

Our systematic assessment and analysis of the AlphaFold database’s protein structure predictions with regard to non-trivial topologies has significantly advanced our understanding of the diversity and conservation of knotted protein families. Using this data, we constructed a large-scale deep learning model that successfully classifies proteins as either knotted or unknotted based on their amino acid sequences. The exceptional accuracy and resilience of our model showcase its potency in discerning sequence characteristics critical for the formation of complex topologies. This achievement not only validates the utility of deep learning techniques in enriching our understanding and promoting research in computational proteomics, but also underscores the promise of artificial intelligence in advancing scientific discovery.

Furthermore, our application of interpretive techniques has illuminated specific patterns associated with conserved knots in SPOUT family proteins. These patterns are particularly pronounced in the areas of the cofactor binding site, a region crucial to the enzymatic function of these proteins. Our findings underscore the importance of this knot in maintaining protein function and raise interesting questions about the role of knotted structures in other protein families.

However, there remains an extensive scope for further inquiry. The search for additional sequence patterns in other knotted protein families presents an exciting avenue for future investigation. This will not only further enhance our understanding of the role of protein knots in biological functions, but will also offer a deeper insight into the profound potential of cutting-edge artificial intelligence technologies in the field of computational proteomics.

## Models and Methods

### Dataset

#### Knot detection in AlphaFold database

With the introduction of AlphaFold version 3, which for the first time revealed predictions of all protein structures from the UniProt database (nearly 200 million entries), knot detection and identification entered the Big Data age. Despite the relative ease of detecting a knot in a single structure using the Topoly Python package, ^25^ the computational demands scale with the data size. Therefore, to address the needs and manage the size of computations an asynchronous distributed task management solution — the kafka-slurm-agent^36^ was implemented earlier and for the purpose of this study significantly enhanced.

The knot detection process was divided into two stages. In the initial stage, protein structures were grouped into batches of 4,000. Structures with a pLDDT score below 70 were discarded, and for the remaining, the HOMFLY-PT polynomial was computed using 100 random closures with a maximum of 60 crossings, as described in. ^26,27^ This stage took approximately two weeks, with the kafka-slurm-agent’s cluster agents distributing the batches over three Linux clusters, collectively consisting of more than 70 nodes and approximately 2,000 cores. Approximately 700,000 structures with an unknot probability less than 0.5 were selected for the next stage.

Stage 2 involved running the HOMFLY-PT polynomial with 500 random closures. Additional metadata from UniProt, such as taxonomy, PFAM, and InterPro annotations, were obtained. The position of the knot core was determined using a fast heuristic algorithm, adapted from the one developed for the AlphaKnot database. ^4^ In light of the clusters’ unavailability, most of this stage was computed on ten individual workstations managed by worker-agents from the kafka-slurm-agent package. Structures with an unknot probability remaining lower than 0.5 and the probability of a particular knot exceeding 0.4 were classified as knotted.

This two-stage knot detection procedure was reapplied to the 9% of structures updated in AlphaFold version 4.

#### Selection of protein families

Based on the resulting dataset of proteins with a detected knot in the predicted AlphaFold structure, proteins were categorised with respect to their assigned InterPro ^37^ identifiers (describing structural domains, families and homologous superfamilies). Minimum subsets of such collections of identifiers were then selected, i.e. the sets of the aforementioned structural feature identifiers whose representatives further manifested a non-trivial topology, in order to filter out redundant IDs. The dataset so obtained was then subjected to quantitative search followed by manual verification, in which the most numerous groups (above 500 proteins) containing examples of proteins with experimentally resolved structures possessing a knot (based on the KnotProt database^5,38,39^ and groups of proteins with either close homology to experimentally confirmed knotted ones or new groups plausible to be knotted (as of the significant amount of non-trivial topologies) were chosen. This ultimately produced a list of 20 different combinations of InterPro identifiers, included in Table 1. Due to the presence of a single domain in several different families, we found duplicate proteins in our dataset (Supplementary Table 2). However, in further steps, we perform sequence clustering, eliminating the repeated protein sequences.

For each set of InterPro identifiers, we compiled a list of UniProt identifiers for the associated proteins. We then created a knotted dataset of protein sequences from the original AlphaFold calculation. We also created an unknotted dataset in a similar manner using the same collection of UniProt IDs, but we excluded the structures identified as potentially knotted in the first stage of knot detection to ensure accurate data labeling. Most of the data manipulation and processing were performed in-house utilizing an installation of Apache Spark and Hadoop.

#### Data preprocessing

To mitigate redundancy and prevent data leakage between the training and testing datasets, we adopted CD-HIT (Cluster Database at High Identity with Tolerance, ^40^) for clustering our protein sequence data prior to the training of the machine learning model. The CD-HIT was configured with a sequence identity threshold of 90%, ensuring sequences sharing a local identity at or above this threshold were categorized into the same cluster. The minimum alignment coverage for the shorter sequences was set at 90%, and the maximum sequence length discrepancy was fixed at 80% of the length of the longer sequences. This ensured the inclusion of only those sequences with substantial alignment coverage. Following this, we chose a single representative sequence from each cluster.

Given the discrepancies in length distributions between knotted and unknotted protein sequences, there was a risk that neural network models could unintentionally rely on sequence length as a classification feature. To address this concern, the datasets were subjected to downsampling, ensuring a balanced sequence length distribution between both protein sequence datasets.

he final dataset comprised 98528 knotted proteins and an equivalent number of unknotted proteins. This was subsequently randomly partitioned into training and testing datasets, with 157644 (80%) samples designated for training and 39412 samples (20%) set aside for testing.

### Neural network models

Our primary objective was to engineer machine learning models capable of classifying knotted proteins based solely on their amino acid sequence.

The best-performing model was based on the ProtBert-BFD model architecture, ^30^ a derivative of the BERT language model that has been pretrained on a vast corpus of amino acid sequences, allowing it to capture important biophysical properties of the proteins.

The model and the tokenizer were downloaded from HuggingFace Hub `Rostlab/prot bert bfd` repository. To determine the optimal parameters for training, we performed a hyperparam-eter search in relation to `gradient accumulation`, `weight decay`, and `learning rate`. The training set initially described was subsequently divided, with 90% of the sequences earmarked for training and the remaining 10% used for comparing models trained with different sets of hyperparameters. The tuning with the best parameters resulted in a `gradient accumulation = 16`, `weight decay = 0`, and `learning rate = 1e-5`. Given the size of the dataset and acceptable convergence, we run only one training epoch. The model training required approximately 2.5 hours on a single A100 GPU.

In addition to fine-tuning 420 million trainable parameters of a large BERT-like neural network model (M1), we also explored two alternate strategies: Employing protein-level embeddings from the same model (420M fixed parameters) and training a smaller neural network (16M trainable parameters) on top of them (M2), and training a small convolutional neural network (M3) on a one-hot encoded sequence (10k trainable parameters). Refer to the Supplement for a detailed description of models M2 and M3.

The models were implemented in Python using either Pytorch^41^ (for M1, M2) or TensorFlow’s Keras^42^ (for M3). We leveraged the MetaCentrum computation cluster, which featured hosted JupyterLab notebooks and A10/A40/A100 GPUs, Python 3.8, PyTorch 1.13, TensorFlow 2.11, and HuggingFace transformers 4.24. For the code and the links to the trained models on Hugging Face hub, please visit our GitHub repository: https://github.com/ML-Bioinfo-CEITEC/pknots_experiments

All models were assessed employing standard accuracy metrics. To examine how the models performed across different protein families, and to ensure the models weren’t solely recognizing a specific subset of families, we also calculated and monitored accuracy per family, true positive rate (TPR), and true negative rate (TNR).

### Interpretation with patching technique

The core problem we addressed was binary classification of proteins as either knotted or unknotted. The secondary aim was to identify the crutial segments of protein sequences that largely contribute to the formation of a knot. Given the availability of the actual knot core locations within knotted sequences, we conducted an evaluation to measure the model’s capability to not only correctly classify an input sequence, but also to pinpoint the knot core’s location. However, conventional interpretability techniques, like Layer Integrated Gradients,^43^ have limited applicability on biological data since they can only relate to individual input points (in our case amino acids), whereas protein folding requires the cooperation of groups of them simultaneously. Therefore, we proposed a patching technique: we monitored how the model’s prediction changed after replacing a continuous segment of amino acids in the original sequence with *X* characters, with respect to the model’s prediction of the original unpatched sequence. Where the *X* character was chosen because it is present in the ProtBert-BFD tokenizer and it signifies an arbitrary amino acid, and the patch size was set specifically for each protein family based on their average knot core and sequence lengths.

Our hypothesis is that if we patch a part of the sequence corresponding to the knot core, the prediction score will drop (from knotted to unknotted), reflecting the significance of such segment. In practical terms, if the prediction score fell below 0.5, the corresponding patch was deemed a candidate for the knot core location. For each sequence we generated its patched versions by moving the patch from left to right with stride 1, and then fed them to the model to obtain their predictions. The patch that resulted in the overall minimum prediction score among all patched versions was chosen as the predicted knot core location. This approach was evaluated by calculating the overlap of the minimum patch location with the actual knot core. Supplementary Figure 2 illustrates this technique for one input sequence.

## Acknowledgement

This work was supported by the National Science Centre (#UMO-2018/31/B/NZ1/04016 and 2021/43/I/NZ1/03341 to JIS). PS, DS and EK were supported by the OPUS LAP program of the Grant Agency of Czech Republic (Reg. No. 204/07/1592 grant “Biological code of knots – identification of knotted patterns in biomolecules via AI approach”). Computational resources were supplied by the project “e-Infrastruktura CZ” (e-INFRA CZ LM2018140) supported by the Ministry of Education, Youth and Sports of the Czech Republic.

## References

(1) Takusagawa, F.; Kamitori, S. A Real Knot in Protein. Journal of the American Chemical Society 1996, 118, 8945–8946.

(2) Mansfield, M. Are there knots in proteins? Nature structural biology 1994, 1, 213–4.

(3) Dabrowski-Tumanski, P.; Rubach, P.; Goundaroulis, D.; Dorier, J.; Sulkowski, P.; Millett, K. C.; Rawdon, E. J.; Stasiak, A.; Sulkowska, J. I. KnotProt 2.0: a database of proteins with knots and other entangled structures. Nucleic acids research 2019, 47, D367–D375.

(4) Niemyska, W.; Rubach, P.; Gren, B. A.; Nguyen, M.; Garstka, W.; Bruno da Silva, F.; Rawdon, E.; Sulkowska, J. AlphaKnot: server to analyze entanglement in structures predicted by AlphaFold methods. Nucleic Acids Research 2022, 50, W44–W50.

(5) Su-lkowska, J. I.; Rawdon, E. J.; Millett, K. C.; Onuchic, J. N.; Stasiak, A. Conservation of complex knotting and slipknotting patterns in proteins. Proceedings of the National Academy of Sciences 2012, 109, E1715–E1723.

(6) Jackson, S. E.; Suma, A.; Micheletti, C. How to fold intricately: using theory and experiments to unravel the properties of knotted proteins. Current Opinion in Structural Biology 2017, 42, 6–14, Folding and binding • Proteins: Bridging theory and experiment.

(7) Tkaczuk, K.; Dunin-Horkawicz, S.; Purta, E.; Bujnicki, J. Structural and evolutionary bioinformatics of the SPOUT superfamily of methyltransferases. BMC bioinformatics 2007, 8, 73.

(8) Christian, T.; Sakaguchi, R.; Perlinska, A. P.; Lahoud, G.; Ito, T.; Taylor, E. A.; Yokoyama, S.; Sulkowska, J. I.; Hou, Y.-M. Methyl transfer by substrate signaling from a knotted protein fold. Nature Structural &amp Molecular Biology 2016, 23, 941–948.

(9) Zayats V, J. A. J. B. D.-H. S. S. J., Perlinska AP Slipknotted and unknotted monovalent cation-proton antiporters evolved from a common ancestor. PLoS computational biology 2021, 8.

(10) Potestio, R.; Micheletti, C.; Orland, H. Knotted vs. unknotted proteins: evidence of knot-promoting loops. PLoS computational biology 2010, 6, e1000864.

(11) Wallin, S.; Zeldovich, K. B.; Shakhnovich, E. I. The Folding Mechanics of a Knotted Protein. Journal of Molecular Biology 2007, 368, 884–893.

(12) Sulkowska, J. I.; Sulkowski, P.; Onuchic, J. Dodging the crisis of folding proteins with knots. Proceedings of the National Academy of Sciences of the United States of America 2009, 106, 3119–24.

(13) King, N. P.; Jacobitz, A. W.; Sawaya, M. R.; Goldschmidt, L.; Yeates, T. O. Structure and folding of a designed knotted protein. Proceedings of the National Academy of Sciences 2010, 107, 20732–20737.

(14) Bustamante, A.; Sotelo-Campos, J.; Guerra, D.; Floor, M.; Wilson, C.; Bustamante, C.; Baez, M. The energy cost of polypeptide knot formation and its folding consequences. Nature Communications 2017, 8.

(15) Joanna, S.; Piotr, S.; Piotr, S.; Marek, C. Stabilizing effect of knots on proteins. Proceedings of the National Academy of Sciences of the United States of America 2009, 105, 19714–9.

(16) Varadi, M.; Anyango, S.; Deshpande, M.; Nair, S.; Natassia, C.; Yordanova, G.; Yuan, D.; Stroe, O.; Wood, G.; Laydon, A., et al. AlphaFold Protein Structure Database: massively expanding the structural coverage of protein-sequence space with high-accuracy models. Nucleic acids research 2022, 50, D439–D444.

(17) Jumper, J.; Evans, R.; Pritzel, A.; Green, T.; Figurnov, M.; Ronneberger, O.; Tunyasuvunakool, K.; Bates, R.; Žídek, A.; Potapenko, A., et al. Highly accurate protein structure prediction with AlphaFold. Nature 2021, 596, 583–589.

(18) Ferrario, E.; Miggiano, R.; Rizzi, M.; Ferraris, D. M. The integration of AlphaFold-predicted and crystal structures of human trans-3-hydroxy-l-proline dehydratase reveals a regulatory catalytic mechanism. Computational and Structural Biotechnology Journal 2022, 20, 3874–3883.

(19) da Silva, F. B.; Lewandowska, I.; Kluza, A.; Niewieczerzal, S.; Augustyniak, R.; Sulkowska, J. I. First crystal structure of double knotted protein TrmD-Tm1570 – inside from degradation perspective. bioRxiv 2023,

(20) Brems, M. A.; Runkel, R.; Yeates, T. O.; Virnau, P. AlphaFold predicts the most complex protein knot and composite protein knots. Protein Science 2022, 31.

(21) Perlinska, A. P.; Niemyska, W. H.; Gren, B. A.; Bukowicki, M.; Nowakowski, S.; Rubach, P.; Sulkowska, J. I. scpAlphaFold/scp predicts novel human proteins with knots. Protein Science 2023, 32.

(22) Andersson, F. I.; Pina, D. G.; Mallam, A. L.; Blaser, G.; Jackson, S. E. Untangling the folding mechanism of the 52-knotted protein UCH-L3. The FEBS journal 2009, 276, 2625–2635.

(23) Virnau, P.; Mirny, L. A.; Kardar, M. Intricate knots in proteins: Function and evolution. PLoS computational biology 2006, 2, e122.

(24) Strassler, S. E.; Bowles, I. E.; Dey, D.; Jackman, J. E.; Conn, G. L. Tied up in knots: Untangling substrate recognition by the SPOUT methyltransferases. Journal of Biological Chemistry 2022, 298.

(25) Dabrowski-Tumanski, P.; Rubach, P.; Niemyska, W.; Gren, B. A.; Sulkowska, J. I. Topoly: Python package to analyze topology of polymers. Briefings in Bioinformatics 2020, 22, bbaa196.

(26) Freyd, P.; Yetter, D.; Hoste, J.; Lickorish, W.; Millett, K.; Ocneanu, A. A new polynomial invariant of knots and links. Bull. Amer. Math. Soc. 1985, 12.

(27) Przytycki, J. H.; Traczyk, P. Invariants of links of Conway type. 2016,

(28) Jarmolinska, A. I.; Perlinska, A. P.; Runkel, R.; Trefz, B.; Ginn, H. M.; Virnau, P.; Sulkowska, J. I. Proteins’ knotty problems. Journal of Molecular Biology 2019, 431, 244–257.

(29) Sikora, M.; Flapan, E.; Wong, H.; Rubach, P.; Garstka, W.; Niewieczerzal, S.; Rawdon, E. J.; Sulkowska, J. I. Proteins containing 6-crossing knot types and their folding pathways. bioRxiv 2023, 2023–06.

(30) Elnaggar, A.; Heinzinger, M.; Dallago, C.; Rehawi, G.; Wang, Y.; Jones, L.; Gibbs, T.; Feher, T.; Angerer, C.; Steinegger, M.; Bhowmik, D.; Rost, B. ProtTrans: Toward Understanding the Language of Life Through Self-Supervised Learning. IEEE Transactions on Pattern Analysis and Machine Intelligence 2022, 44, 7112–7127.

(31) Keren, H.; Lev-Maor, G.; Ast, G. Alternative splicing and evolution: diversification, exon definition and function. Nature Reviews Genetics 2010, 11, 345–355.

(32) Zayats, V.; Perlinska, A. P.; Jarmolinska, A. I.; Jastrzebski, B.; Dunin-Horkawicz, S.; Sulkowska, J. I. Slipknotted and unknotted monovalent cation-proton antiporters evolved from a common ancestor. PLoS Computational Biology 2021, 17, e1009502.

(33) Shi, D.; Allewell, N. M.; Tuchman, M. From genome to structure and back again: A family portrait of the transcarbamylases. International Journal of Molecular Sciences 2015, 16.

(34) Perlinska, A. P.; Stasiulewicz, A.; Nawrocka, E. K.; Kazimierczuk, K.; Setny, P.; Sulkowska, J. I. Restriction of S-adenosylmethionine conformational freedom by knotted protein binding sites. PLoS computational biology 2020, 16, e1007904.

(35) Lundberg, S. M.; Lee, S.-I. A unified approach to interpreting model predictions. Advances in neural information processing systems 2017, 30.

(36) https://github.com/prubach/kafka-slurm-agent.

(37) Paysan-Lafosse, T. et al. InterPro in 2022. Nucleic Acids Research 2022, 51, D418–D427.

(38) Dabrowski-Tumanski, P.; Rubach, P.; Goundaroulis, D.; Dorier, J.; Su-lkowski, P.; Millett, K. C.; Rawdon, E. J.; Stasiak, A.; Sulkowska, J. I. KnotProt 2.0: a database of proteins with knots and other entangled structures. Nucleic Acids Research 2018, 47, D367–D375.

(39) Jamroz, M.; Niemyska, W.; Rawdon, E. J.; Stasiak, A.; Millett, K. C.; Su-lkowski, P.; Sulkowska, J. I. KnotProt: a database of proteins with knots and slipknots. Nucleic Acids Research 2014, 43, D306–D314.

(40) Fu, L.; Niu, B.; Zhu, Z.; Wu, S.; Li, W. CD-HIT: accelerated for clustering the nextgeneration sequencing data. Bioinformatics 2012, 28, 3150–3152.

(41) Paszke, A. et al. Advances in Neural Information Processing Systems 32 ; Curran Associates, Inc., 2019; pp 8024–8035.

(42) Chollet, F., et al. Keras. https://keras.io,2015.

(43) Cik, I.; Rasamoelina, A. D.; Mach, M.; Sincak, P. Explaining Deep Neural Network using Layer-wise Relevance Propagation and Integrated Gradients. 2021.

