## Supplementary Material for "Knot or Not? Sequence-Based Identification of Knotted Proteins With Machine Learning"

Denisa Šrámková*, Maciej Sikora*, Dawid Uchal, Eva Klimentová, Agata P. Perlinska, Mai Lan Nguyen, Marta Korpacz, Roksana Malinowska, Pawel Rubach, Petr Šimeček, Joanna I. Sulkowska

#### M2 model

The M2 model is a simple convolutional neural network (CNN) that leverages protein embeddings from the ProtBert-BFD model [1]. These embeddings are processed using average pooling to yield a fixed-size vector of 1024 dimensions. The CNN is composed of a convolutional layer with a kernel size of 7 and 32 channels, followed by batch normalization and a ReLU activation function. The output is then flattened and passed through two linear layers before being processed by a single neuron with a sigmoid activation function, yielding a probability of the input sequence being knotted.

The model was trained over 10 epochs using Adam optimizer with default settings, learning rate 0.0001 and a batch of size 32. The model reached 96.9 % accuracy on the test set.

#### M3 model

The M3 model, a CNN, is less complex compared to the other two, borrows its architecture from the PENGUINN model [2]. We selected training and testing data such that sequences exceeding 500 amino acids were filtered out. Sequences shorter than 500 amino acids were padded with the 'X' character to reach the uniform length of 500. The sequences were then one-hot encoded to serve as inputs to the model.

Training of the M3 model spanned over 10 epochs using the Adam optimizer with default parameters, a batch size of 128, and a validation split of 0.3. Upon evaluation of the test set, it achieved an accuracy of 95.18%.

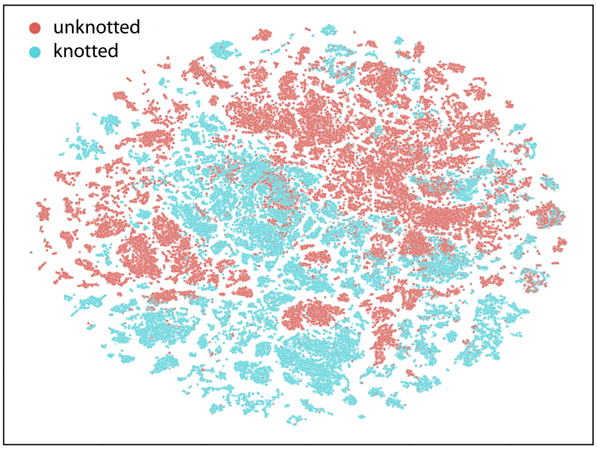


Figure 1: Visualization of protein sequences embeddings from ProtBert-BFD using t-SNE technique, coloured according to topology type in the training set.

#### Validation on ketol-acid reductoisomerases

For independent validation, we selected a family of ketol-acid reductoisomerases, excluded from the training set. Experimental verifications suggest that some proteins in this family possess a $4_1$ knot [3]. The sequences were processed similarly to the main dataset (clustering, train/test split set at an 80:20 ratio). Totally, we have 2304 proteins in the training set (794 knotted, 1510 unknotted) and 577 proteins in the test set (225 knotted, 352 unknotted).

Initially, we tested the pre-trained models M1, M2, and M3. As depicted, the accuracy of the predictions generally suffers for either positive or negative cases. This discrepancy likely stems from

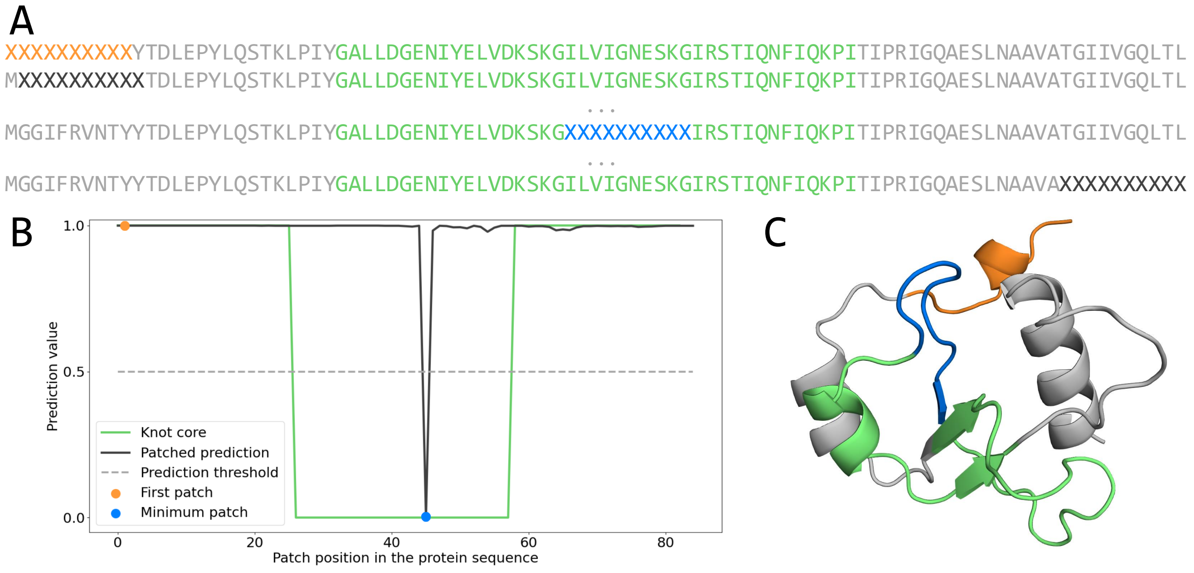


Figure 2: **The patching technique demonstrated on one SPOUT protein (UniProtKB ID: A0A2W5F4Z7).** A. The patched versions of the input sequence with two specific patches highlighted (first: orange, minimum: blue). B. A plot of the prediction scores for all patched versions of the sequence. We can observe one significant drop below the 0.5 (knotted to unknotted) threshold and its corresponding location (blue). C. The 3D protein structure with highlighted knot core (green), minimum patch (blue), and the first patch (orange).

Table 1: **Evaluation of models on ketol-acid reductoisomerases family.** False positive rate (FPR) and false negative rate (FNR) of six models evaluated on the test set of the ketol-acid reductoisomerases family.

| Model | FPR (%) | FNR (%) |
|---|---|---|
| M1 | 57.94 | 8.44 |
| M2 | 67.61 | 2.22 |
| M3 | 34.64 | 77.22 |
| M1b | 8.24 | 1.33 |
| M2b | 2.84 | 8.00 |
| M3b | 3.88 | 6.74 |

family-specific knotting features that the models had not been trained to detect. We can conclude, that the model cannot be relied on to predict the knots outside its domain (i.e. families it has been trained on). This discrepancy can be visualized as a T-SNE plot of protein embeddings, where predictions from M2 are on the left, and the true labels (knotted vs unknotted) are on the right, as shown in Supplementary Figure 3.

Not only do we observe that the current models are not performing well, but it is also evident the knotted and unknotted cases in Supplementary Figure 3 can be separated. For that, we fine-tuned M1, M2, and M3 specifically for this family, resulting in models M1b, M2b, and M3b. A comparison of false positive rates across all six models is displayed in Supplementary Table 1.

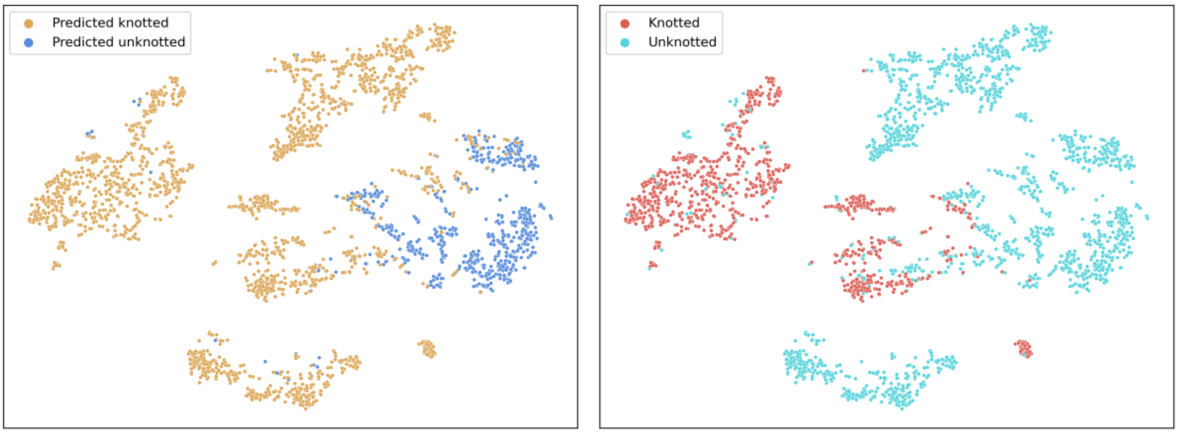


Figure 3: Visualization of protein sequence ProtBERT-BFD embeddings from TODO family using t-SNE technique. The left panel is coloured according to M2 model predictions, the right panel is coloured based on true topology.

Table 2: **Overlap between IPR002035 (von Willebrand factor, type A) and other families/domains in our dataset.** Protein numbers are based on the UniProtKB.

| Family ID | Family/domain name | No. proteins with both domains |
|---|---|---|
| IPR013694 | VIT | 18562 |
| IPR032695 | Integrin domain superfamily | 6774 |
| IPR013649 | Integrin alpha-2 | 6318 |
| IPR036398 | Alpha carbonic anhydrase domain superfamily | 15 |
| IPR004837 | Sodium/calcium exchanger membrane region | 2 |
| IPR029026 | tRNA (guanine-N1-)-methyltransferase, N-terminal | 1 |

### References


[1] A. Elnaggar, M. Heinzinger, C. Dallago, G. Rehawi, Y. Wang, L. Jones, T. Gibbs, T. Feher, C. Angerer, M. Steinegger, D. Bhowmik, and B. Rost, "Prottrans: Toward understanding the language of life through self-supervised learning," *IEEE Transactions on Pattern Analysis and Machine Intelligence*, vol. 44, no. 10, pp. 7112–7127, 2022.

[2] E. Klimentova, J. Polacek, P. Simecek, and P. Alexiou, "PENGUINN: Precise exploration of nuclear g-quadruplexes using interpretable neural networks," *Front. Genet.*, vol. 11, p. 568546, Oct. 2020.

[3] P. Virnau, L. A. Mirny, and M. Kardar, "Intricate knots in proteins: Function and evolution," *PLoS computational biology*, vol. 2, no. 9, p. e122, 2006.
